# Quantifying Codon Usage in Signal Peptides: Gene Expression and Amino Acid Usage Explain Apparent Selection for Inefficient Codons

**DOI:** 10.1101/347849

**Authors:** Alexander L. Cope, Robert L. Hettich, Michael A. Gilchrist

## Abstract

The Sec secretion pathway is found across all domains of life. A critical feature of Sec secreted proteins is the signal peptide, a short peptide with distinct physicochemical properties located at the N-terminus of the protein. Previous work indicates signal peptides are biased towards translationally inefficient codons, which is hypothesized to be an adaptation driven by selection to improve the efficacy and efficiency of the protein secretion mechanisms. We investigate codon usage in the signal peptides of *E. coli* using the Codon Adaptation Index (CAI), the tRNA Adaptation Index (tAI), and the ribosomal overhead cost formulation of the stochastic evolutionary model of protein production rates (ROC-SEMPPR). Comparisons between signal peptides and 5’-end of cytoplasmic proteins using CAI and tAI are consistent with a preference for inefficient codons in signal peptides. Simulations reveal these differences are due to amino acid usage and gene expression - we find these differences disappear when accounting for both factors. In contrast, ROC-SEMPPR, a mechanistic population genetics model capable of separating the effects of selection and mutation bias, shows codon usage bias (CUB) of the signal peptides is indistinguishable from the 5’-ends of cytoplasmic proteins. Additionally, we find CUB at the 5’-ends is weaker than later segments of the gene. Results illustrate the value in using models grounded in population genetics to interpret genetic data. We show failure to account for mutation bias and the effects of gene expression on the efficacy of selection against translation inefficiency can lead to a misinterpretation of codon usage patterns.

## Introduction

A secreted protein can broadly be defined as any protein entering a secretory pathway for transport through a cellular membrane. These proteins serve important cellular functions, including metabolism and antibiotic resistance [15, 37]. Secreted proteins also play essential roles in the virulence of pathogenic bacteria [15]. Numerous secretion systems exists and vary between and within taxa [1, 15, 37]. Despite the diversity of secretion pathways, the general secretion pathway, also commonly referred to as the Sec pathway, is found across all domains of life [15, 26]. In brief, proteins are transported to the SecYEG translocon located in the membrane in a chaperone-dependent (SecA/B and SRP) or chaperone-independent manner [26, 43]. All SecA/B-dependent proteins and chaperone-independent, as well as some SRP-dependent proteins, contain a short peptide chain located at the N-terminus of the protein known as the signal peptide [15, 26, 43]. The signal peptide is an essential component of the Sec pathway, serving as a binding site for the appropriate chaperones and/or helping delay the folding of the protein [26, 43]. Although signal peptides do vary in their amino acid sequences, signal peptides have distinct physicochemical properties which biases their amino acid usage [26, 43, 49]. A signal peptide generally consists of 3 regions: a positively charged N-terminus, a hydrophobic core, and a polar C-terminus, where the signal peptide is cleaved from the rest of the protein, sometimes referred to as the ‘‘mature peptide.”

The ability to accurately predict signal peptides is useful for identifying secreted proteins in non-model organisms; this has led to the development of machine learning approaches to predict signal peptides which take advantage of the distinct physicochemical properties of signal peptides, such as SignalP [31]. Although the physicochemical properties of signal peptides are consistent, altering the N-terminus has a range of effects on protein secretion: from a decrease in the number of proteins secreted to no observable effect [18, 27, 34, 45]. The variability in the outcomes of neutralizing the N-terminal positive charge led to a search for other mechanisms which also contribute to the efficacy of protein secretion [49, 50].

Numerous studies suggests codon usage bias (CUB) - the non-uniform usage of synonymous codons - contributes to effective protein secretion in *E. coli* [3, 32, 52, 51, 53, 55]. found *E. coli* K12 MG1655 signal peptides are biased for translation inefficient codons, which are predicted to be translated slower than their synonymous counterparts. This is in stark contrast to the rest of the *E. coli* proteome, where *E. coli* is biased towards the most efficient codons [17, 32]. [20, 21, 24] examined the usage of inefficient codons in signal peptides of *S. coelicolor, S. cerevisiae*, and various multicellular eukaryotes and came to similar conclusions when applying codon usage indices such as the Codon Adaptation Index (CAI) [41] and tRNA Adaptation Index (tAI) [7]. Consistent across this work is the interpretation that selection is driving the apparent increase in inefficient codon usage in signal peptides. Furthermore, [54] concluded an overabundance of the lysine codon AAA at the second position in the signal peptide promoted efficient translation initiation.

[49] hypothesized an adaptive role for inefficient codons in the protein secretion process in which the combination of efficient translation initiation and inefficient translation reduced the distance between sequential ribosomes along the mRNA, leading to more efficient recycling of the necessary chaperones. Other explanations for the observed increase in inefficient codons include the inability of *E. coli* SRP to induce a translational pause following signal peptide recognition [33, 49] and slowing down the co-translational folding of the protein, as a folded protein cannot be translocated through the SecYEG translocon [32, 52, 51, 50]. If signal peptides have a different CUB relative to the rest of the genome, then codon-level information could be incorporated into signal peptide prediction tools.

In contrast [21] found no significant differences in the ribosome densities between the signal peptides and the 5’-ends of nonsecretory genes in various eukaryotes. Ribosome densities are expected to be higher in signal peptides relative to the 5’-end of nonsecretory genes if selection is acting to increase translation inefficiency in the signal peptide. Additionally, while both [24] and [21] examined codon usage in relation to secretion in *H. sapiens* using a metric based on tAI, only [24] found results consistent with increased frequencies of inefficient codons in signal peptides. From a population genetics perspective, it is surprising statistically significant results were obtained in a mammal, which usually have little adaptive CUB due to their lower effective population sizes [5, 22]. More recently, [38] found codon optimization of a signal peptide improved localization of the protein to the periplasm of *E. coli*, seemingly contradicting a general role for inefficient codon usage in signal peptides. A potential reason for these contradictions is the previous analyses of signal peptide codon usage by [20, 21, 24, 32] did not adequately account for the effects of mutation bias and drift in shaping codon usage [2, 13, 11, 12, 40, 46].

We re-examined CUB in signal peptides of *E. coli* using CAI, tAI, and ROC-SEMPPR a population genetics model which accounts for selection, mutation bias, and gene expression - to determine if selection on codon usage in signal peptides differs from the 5’-ends of genes. Although we find significant differences in codon usage using CAI and tAI, we present evidence these differences are due to signal peptide-specific amino acid biases and differences in the gene expression distributions of genes with and without signal peptides. When comparing signal peptides and the 5’-ends of genes not containing a signal peptide with ROC-SEMPPR, we find signal peptide codon usage is consistent with the 5’-ends. We find selection on codon usage favors the efficient codons, but the strength of selection is weaker at the 5’-ends, corroborating previous analyses [9, 13, 11, 32, 35].

Our work demonstrates the value of analyzing CUB from a formal population genetics framework, as well as highlights potential limitations with using more common metrics such as CAI for analyzing codon usage on relatively small regions of the genome. Failure to account for variation in the strength of selection due to variation in gene expression can lead to conflating mutation bias with selection, resulting in a misinterpretation of observed codon usage patterns. Our work also illustrates the importance of considering non-adaptive forces in shaping biological phenomenon before invoking adaptive explanations [14]. We believe this is particularly important in the modern genomic-age when the combination of large datasets, misinterpretation of p-values, and an inherent bias towards adaptationist interpretations can lead to the proliferation of over-interpreted hypotheses within the biological community.

## Materials and Methods

### Signal Peptide Prediction

Signal peptides were predicted using SignalP 4.1 [31] using both the default cutoff D-score of 0.51 and a more conservative D-score of 0.75. In brief, SignalP consists of two neural networks, one for determining the amino acid sequence similarity to signal peptides and the other for identifying the most likely cleavage site. The results of both neural networks are combined into one value, called the D-score, which ranges between 0 and 1. Setting the cutoff D-score closer to 1 results in a lower false positive rate. A set of confirmed signal peptides for *E. coli* K12 MG1655 was taken from The Signal Peptide Website (http://www.signalpeptide.de/). All analyses in the main text will focus on the set of signal peptides with *D ≥* 0.51 as this set provides us with the most data; analyses of the *D* > 0.75 and set of confirmed signal peptides give similar results (see Supplementary Material).

### ROC-SEMPPR

Given a set of protein-coding genes, ROC-SEMPPR employs a Markov Chain Monte Carlo (MCMC) to estimate codon specific parameters for mutation bias Δ*Μ* and pausing times Δ*η* for each codon within a synonymous codon family (Table 1). In previous work, Δ*η* was scaled relative to the most efficient codon, which had Δ*η* and Δ*Μ* values fixed at 0. To avoid the choice of reference codon affecting our comparisons of CUB between regions, all Δ*η* values in this paper are re-scaled such that these values are centered around 0 for each amino acid. The Δ*η* values reflect the strength and direction of selection against translation inefficiency in a set of protein-coding regions (e.g. the signal peptides). A region with stronger selection against translation inefficiency will have higher Δ*η* values on average than a region with weaker selection. Similarly, a region which favors translation inefficiency would be expected to have Δ*η* values which negatively correlate with a region which favors translation efficiency.

**Table 1:** Description of ROC-SEMPPR parameters used in this paper.

| Parameters | Description |
| --- | --- |
| $\Delta\eta_i$ | Cost of translating codon $i$ relative to reference codon |
| $\Delta M_i$ | Mutation bias towards codon $i$ relative to the reference codon |
| $\phi_k$ | Average Protein Production Rate of gene $k$ |

ROC-SEMPPR also estimates an average protein production rate *φ* for each gene (Table 1). It is important to note ROC-SEMPPR is structured such that the average value of *φ* across the genome is 1. This choice of scaling means the pausing times Δ*η* represent the average strength of selection relative to genetic drift for or against a given codon. We find ROC-SEMPPR estimated *φ* values correlate well with empirical measurements of protein production rates for *E. coli* (see Supplementary Methods: Assessing ROC-SEMPPR Model Adequacy and Figures S1 - S2). If changes in synonymous codon usage alter the efficiency at which a protein is translated, then such a change will have the largest impact on the energetic costs of proteins with high production rates, making *φ* a more appropriate gene expression metric than say, mRNA abundance or protein abundance. Thus, we use protein production rates *φ* as our metric of gene expression. For more details on ROC-SEMPPR, see [12]. Analysis of CUB with ROC-SEMPPR was performed using AnaCoDa [19].

### CAI and tAI

Analysis of CUB was also performed using CAI [41] and tAI [7]. Both CAI and tAI quantify CUB by assigning weights to the 61 sense codons. For CAI, each codon is assigned a weight based on its relative frequency to its synonymous counterparts in a reference set of highly expressed genes, such as ribosomal protein coding genes. The key assumption of CAI is the most frequent codons in the reference set are the most efficient codons [41]. In contrast, tAI assigns weights based on tRNA abundances corresponding to a codon, as well as accounting for codon-anticodon interactions. The key assumption of tAI is the most efficient codons are usually those with the most abundant tRNA [7].

CAI and tAI both range between 0 and 1. A CAI score closer to 1 represents a sequence which more closely resembles the codon usage of the reference set of genes, while a tAI closer to 1 indicates a sequence is more closely adapted to the genomic tRNA pool [7, 41]. Calculations for CAI were performed using the AnaCoDa [19], while tAI was calculated using the R package tAI [6].

### Generating Datasets

Previous analysis of the *E. coli* genome found a set of genes with CAI values that had a negative correlation with their gene expression estimates [8]. It is believed many of these genes were the result of horizontal gene transfer and had not yet reached evolutionary equilibrium with respect to their CUB. We repeated the analysis described in [8] on the current *E. coli* K12 MG1655 genome (version 3, NC_000913.3). Briefly, correspondence analysis was performed using CodonW [30], followed by clustering based on the principle axis scores using the CLARA algorithm [23] in R. Our analysis was consistent with the findings of [8], revealing 782 genes with a CUB deviating significantly from the majority of the *E. coli* genome. We will refer to this set of 782 genes as the “exogenous” component of the genome and the rest of the *E. coli* genome as the “endogenous” for simplicity. All analyses presented will consider only “endogenous” genes because the “exogenous” genes may violate the implicit assumptions of CAI and tAI and the explicit assumptions of ROC-SEMPPR.

Proteins with a signal peptide were split into the signal peptide and the mature peptide the segment of the peptide chain after the signal peptide. On average, the signal peptides were 23 codons long. For comparisons to the 5’-ends of nonsecretory genes - defined here as those lacking a signal peptide - the first 23 codons of the nonsecretory genes were used. We note the secretory genes have an average protein production rate *φ* approximately 10% higher than that of the nonsecretory genes (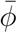 = 1.08 and 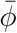 = 0.992,respectively, Figure S3).

As the strength of selection on CUB scales with protein production rate *φ*, we created a control group that eliminates differences in the distribution of *φ* for the nonsecretory genes and signal peptide genes. Specifically, the nonsecretory genes were selected using acceptance-rejection sampling to create the “pseudo-secreted proteins”. In brief, acceptance-rejection sampling is a procedure for sampling from a population such that its distribution of a metric for one population mirrors the distribution of the same metric for another population. In this case, the pseudo-secreted proteins were sampled such that the mean and variance of the log (*φ*) values reflected those of the genes with a signal peptide. The CUB signature of a gene varies with protein production rate *φ*; thus we can be more confident any differences seen between genes with a signal peptide and pseudo-signal peptide genes are not due to differences in their respective *φ* distributions. All pseudo-secreted proteins were split into two regions we will refer to as the “pseudo-signal peptides” and the “pseudo-mature peptides” (the first 23 codons and the remainder of the gene, respectively).

To assess the performance of CAI and tAI when comparing regions with differences in the distributions of protein production rates *φ* and amino acid biases, simulated sequences were used. Sequences based on the 5’-ends of nonsecretory genes, pseudo-signal peptides, and signal peptides were simulated using the AnaCoDa package [19]. To normalize for amino acid usage, sequences 23 amino acids in length were randomly generated to match the amino acid frequencies of the signal peptides. The codon usage of these sequences was also simulated in AnaCoDa, assuming either the *φ* distribution of the nonsecretory genes or the pseudo-secreted proteins. All sequences were simulated using the pausing times Δ*η* and mutation bias Δ*Μ* parameters estimated from the 5’-end of endogenous nonsecretory genes.

### Analysis of Codon Usage with CAI, tAI, and ROC-SEMPPR

We estimated protein production rates *φ* by fitting ROC-SEMPPR to the protein-coding sequences in the *E. coli* K12 MG1655 genome. Analysis of intragenic (e.g. signal vs. mature peptides) and intergenic (e.g. pseudo-signal peptides vs. real signal peptides) CUB was carried out using the mixture distribution functionality available in the AnaCoDa implementation of ROC-SEMPPR [19]. We assumed mutation bias was consistent for the entire genome; thus, we forced mutation bias Δ*Μ* parameters to be equal across the groups of regions. Each group of regions (e.g. signal peptides, mature peptides, etc.) was assumed to have an independent set of pausing time parameters, allowing pausing time Δ*η* estimates to vary between them. *φ* was fixed for each region of a gene at the value estimated when the model was fit to the entire protein-coding sequence. This is done for two reasons: (a) shorter regions, such as the signal peptide, likely have insufficient information to accurately estimate *φ* and (b) this guarantees our gene expression metric has the same impact on the estimates of Δ*η* and Δ*Μ* for intragenic regions, such as a signal peptide and its corresponding mature peptide. We note the use of empirical *φ* estimates in place of ROC-SEMPPR estimated *φ* did not impact our interpretations.

A Model-II regression was used to compare estimated pausing times Δ*η* between regions. Unlike ordinary least squares, Model-II regression, or errors-in-variables regression, accounts for errors in both the *x* and *y* variables [42]. When both variables are subject to error, which is the case for the Δ*η* estimates, the use ordinary least squares leads to downwardly biased parameter estimates. A Model-II regression slope *β* = 1 (or the *y* = *x* line) will serve as the null hypothesis, as this indicates both the strength and direction of selection between two regions are the same. The intercept parameter was fixed at *α* = 0 because the Δ*η* estimates are scaled such that the mean value of Δ*η* is 0. We note that when we allowed the a parameter to vary, it was as expected, approximately 0. For more details on our use of Model-II regression, see Supplementary Methods.

CAI and tAI were used to compare codon usage between signal peptides, 5’-ends, and pseudo-signal peptides [8, 7, 41]. As recommended by [41], methionine and tryptophan were not included when normalizing for the length of the gene in our calculations of CAI. Statistical significance was assessed using a one-tailed Welch’s t-test in R [36]. R and Python scripts used for this paper can be found at https://github.com/acope3/Signal_Peptide_Scripts.

## Results

Our analysis of CUB in signal peptides and the 5’-ends of nonsecretory genes using ROC-SEMPPR revealed these regions to be indistinguishable. Qualitatively, the expected codon frequencies for the 5’-ends of nonsecretory genes and the signal-peptides based on the pausing time Δ*η* and mutation bias Δ*Μ* values estimated from these regions are indistinguishable (Figure S4). Cysteine, aspartic acid, lysine, glutamine, and tyrosine are apparent exceptions, but only the 95% posterior probability intervals of cysteine and glutamine fail to overlap with *y* = *x* line. When comparing the pausing times Δ*η* of signal peptides to the 5’-ends of nonsecretory genes using a Model-II regression, we find no significant difference from the *y* = *x* line (slope *β* 95% confidence interval: 0.923 - 1.128, Figure 1a). To determine if differences were not detected due to underlying differences in the distributions of *φ*, we compared Δ*η* estimated from signal peptides and pseudo-signal peptides. Again, no statistically significant difference from the *y* = *x* line was found and the expected codon frequencies are similar (*β* 95% confidence interval: 0.939 - 1.149, Figure 1b and S5). Similar results are obtained using the signal peptides with a D-score greater than 0.75 or the confirmed signal peptides (Figures S6 - S7). We also see no significant result when using empirically estimated *φ* values (*β* = 0.908, 95% confidence interval: 0.671 - 1.168, Figure S8), although these results show much more variability. The increased variability in the Δ*η* values and corresponding regression line is unsurprising given the empirically estimated *φ* values are subject to significant noise (Figure S2), but are, in this case, treated as error free estimates of a gene’s true *φ* value.

**Figure 1:**
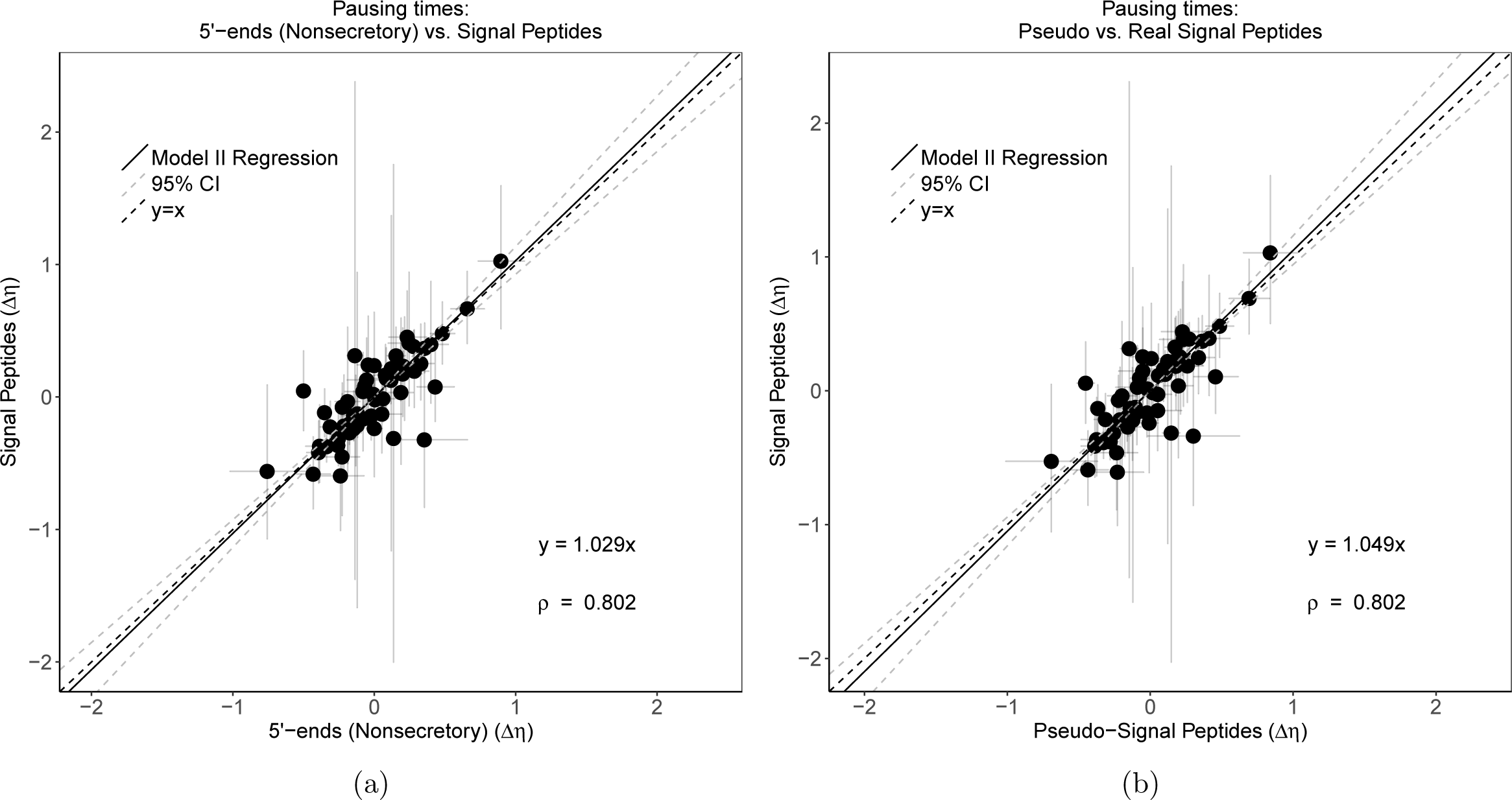
Comparing the pausing time estimates Δ*η* between (a) the 5%-ends of nonsecretory genes or (b) pseudo-signal peptides to signal peptides. Grey dashed lines represent the 95% confidence intervals of the regression line. Results clearly show a strong positive linear relationship (*ρ* = 0.802) between the regions and a regression line not significantly different from *y* = *x*.

The Model-II regression lines estimated from the mature vs. signal peptide comparison and the pseudo-mature vs. pseudo-signal peptide comparison are similar, providing further evidence the nature and magnitude of selection on codon usage in signal peptides and the 5’-ends of nonsecretory genes is indistinguishable (Figure 2). The mature vs. signal peptide comparison produces a regression line with slope *β* = 0.480 (95% confidence interval: 0.428 −0.574), which is approximately 50% of the slope observed when comparing signal peptides to the 5’-ends of nonsecretory genes and pseudo-signal peptides. This indicates selection on codon usage in the mature peptides is stronger than it is in signal peptides, although the nature of selection is still *against* translation inefficiency. Similar behavior is observed when comparing the pseudo-mature vs. pseudo-signal peptide comparison (*β* = 0.509, 95% confidence interval: 0.490 - 0.533). The slope estimate from the mature vs. signal peptide comparison is not significantly different from *β* = 0.509 (Two-tailed Z-test, *p* = 0.0682). Similar regression lines would not be expected if differences in selection on codon usage existed between signal peptides and the pseudo-signal peptides.

**Figure 2:**
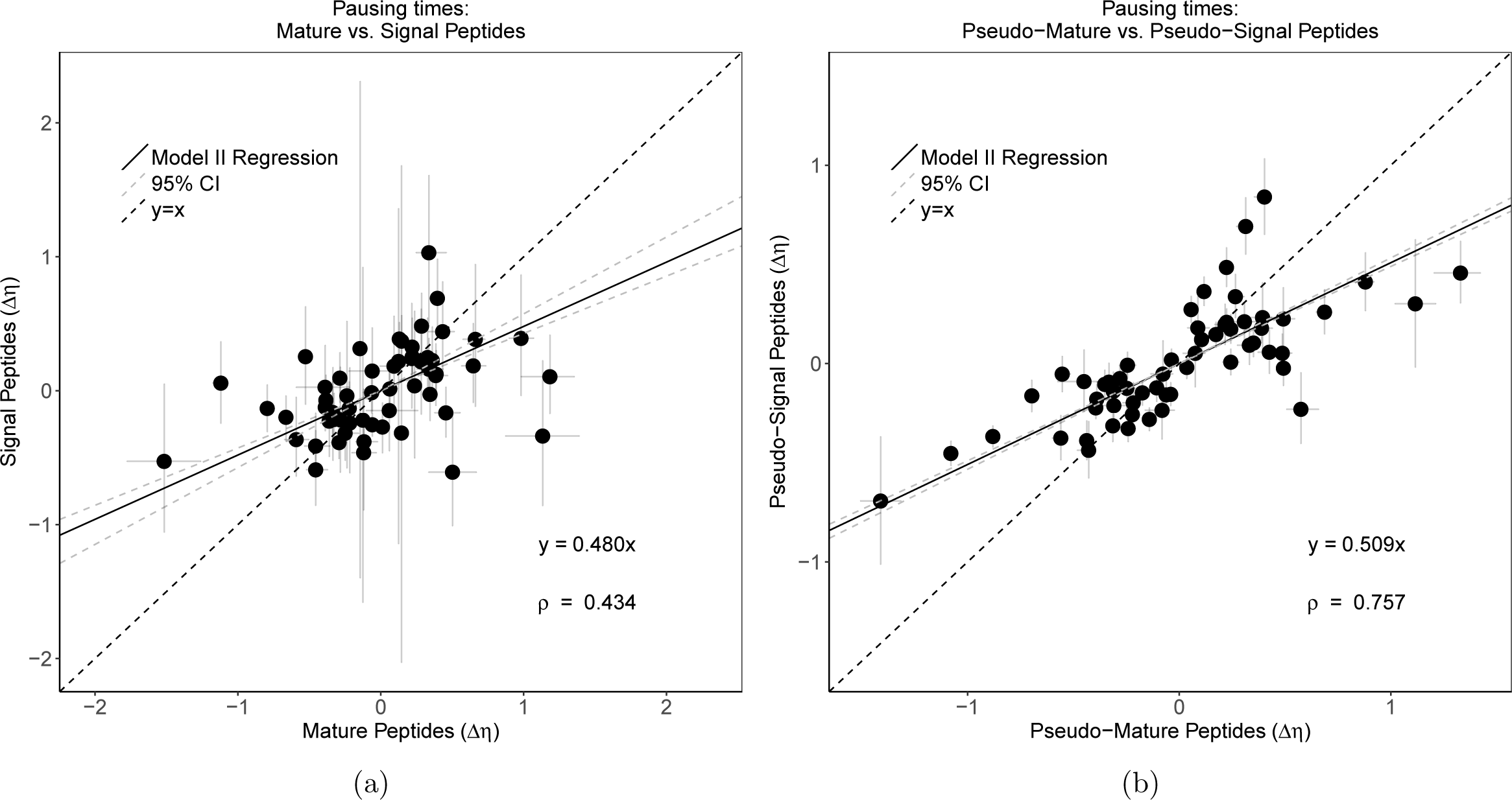
(a) Comparing the codon pausing time estimates Δ*η* between mature peptides and signal peptide regions. Grey dashed lines represent the 95% confidence intervals of the regression line. Results show a positive linear relationship (ρ = 0.43) between the Δ*η* estimates for the two regions. This indicates codons favored in one region tend to be favored in the other. (b) Same comparison for pseudo-signal peptide genes. Regression estimates are indistinguishable from those estimated for the mature and signal peptide comparison (Likelihood Ratio test, *p* = 0.562).

Noting CAI and tAI do not account for the effects of gene expression, mutation bias, drift, or amino acid biases, we found signal peptides have lower CAI and tAI values compared to the first 23 codons of nonsecretory genes (one-tailed Welch’s t-test, *p* < 10^−5^). This was also the case when looking at the pseudo-signal peptides, which normalizes for protein production rates *φ*. These results with CAI and tAI can potentially be explained by either the preferred use of inefficient codons in signal peptides *or* as artifacts of amino acid biases. Signal peptides have a different amino acid composition from the 5’-end due to the required physicochemical properties of this region (Figure S9). We examined the robustness of tAI and CAI as a means of quantifying differences in selection on codon usage when underlying differences between amino acid composition and *φ* exists using data simulated under the same mutation bias Δ*Μ* and pausing time Δ*η* parameters. When comparing simulated signal peptides to simulated 5’-end of nonsecretory genes and simulated pseudo-signal peptides using CAI, the simulated signal peptides are found to have a significantly lower mean CAI (Welch’s t-test, *p* < 0.05) 100% of the time (Figure 3A-B), despite the fact the Δ*η* and Δ*Μ* parameters used to simulate these regions were the same. This suggests differences in amino acid usage and not adaptation to novel selective forces, explains the lower CAI of the signal peptides.

**Figure 3:**
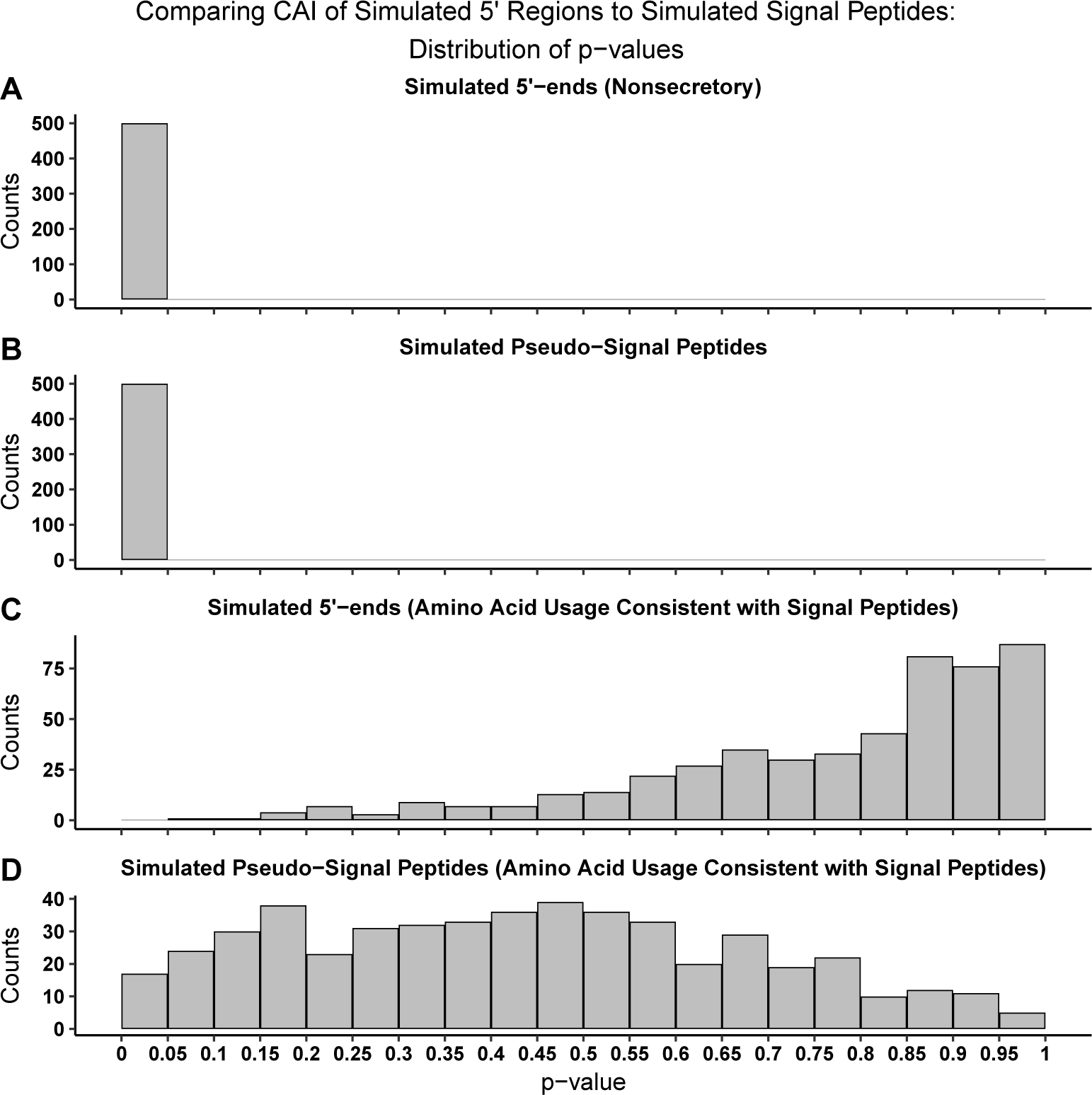
Distribution of p-values from a one-tailed Welch’s t-test comparing CAI in simulated nonsecretory 5’-ends, pseudo-signal peptides, and signal peptides in which all regions were simulated using the same pausing time Δ*η* and Δ*Μ* parameters. (A-B) The CAI of simulated signal peptides was found to be significantly lower on average at a 100% false positive rate when compared to simulated 5’-ends of nonsecretory genes and simulated pseudo-signal peptides. (C) Adjusting the amino acid frequencies of the 5’-end of nonsecretory genes to match those of the signal peptides results in a heavily skewed distribution. (D) Adjusting the amino acid frequencies of the pseudo-signal peptides to match those of the signal peptides results in a more uniform distribution.

When using simulated 5’-ends of nonsecretory genes which have amino acid composition consistent with the signal peptides, the p-values were heavily skewed towards 1. (Figure 3C). This odd behavior is due to the differences in the *φ* distribution differences of the signal peptide and nonsecretory genes. As the former has a higher mean *φ*, the signal peptides on average will have a stronger CUB after normalizing for the amino acid biases. A one-tailed Welch’s t-test with the alternative hypothesis being signal peptides have a lower mean CAI, when in reality they likely have a larger mean CAI, would skew the p-value distribution towards 1. Importantly, ROC-SEMPPR did not detect significant differences between signal peptides and the 5’-ends of non-secretory genes, despite differences in the φ distributions (Figure 1a). When normalizing for both amino acid usage and *φ*, significant differences in CAI are found approximately 4% of the time, which is close to the expected number of false positives at the 0.05 significance level (Figure 3D). Similar results are seen when using tAI (Figure S10). Our results indicate CAI and tAI are prone to inflating differences in CUB between two regions when differences in *φ* and amino acid usage are not accounted for.

Notably, selection on codon usage near the N-terminus appears to be on average approximately 50% weaker than the remainder of the gene based on the slopes *β*. Previous analyses using a variety of codon usage metrics found CUB near the 5’-end to be weaker than middle sections of the gene, with these differences being attributed to selection against nonsense errors and to maintain translation initiation efficiency by reducing mRNA secondary structure [9, 13, 11, 16, 35, 32]. We confirm this trend using ROC-SEMPPR (Figure S11).

[54] proposed selection for translation initiation efficiency was shaping signal peptide codon usage, particularly the use of lysine codon AAA, at the second amino acid position. While AAA appears to be slightly favored in signal peptides, which is not the case in the pseudo-signal peptides, the 95% posterior probability interval overlaps with the *y* = *x* line (Figure S12). If the insignificant increased usage of AAA is due to greater selection for translation initiation efficiency in signal peptides, then removing the first 3 codons when analyzing signal peptide codon usage should remove this effect. Doing so results in no change in the behavior of AAA, suggesting if there is any selection for increased AAA usage in signal peptides, it is not due to selection for increased translation initiation efficiency (Figure S13). Notably, AAA is both mutationally and selectively-favored for lysine in *E. coli.* Keeping in mind selection on CUB is weaker near the 5’-end of the genes in *E. coli*, the combination of weaker selection, mutational favorability, and a slight increase in the occurrence of lysine in signal peptides (Figure S9) likely drives up the frequency of codon AAA in signal peptides relative to the 5’-ends of nonsecretory genes.

## Discussion

In summary, we found no evidence to support the hypothesis that selection on codon usage in signal peptides and the 5’-ends of nonsecretory genes in *E. coli* using a mechanistic model of CUB which incorporates the effects of selection, mutation bias, gene expression, and amino acid usage. We find commonly employed codon usage metrics CAI and tAI produce spurious differences between signal peptides and 5’-ends of nonsecretory genes due to differences in amino acid usage and gene expression of signal peptide containing genes relative to the rest of the genome. Importantly, both amino acid usage and *φ* were significant confounding factors when analyzing CUB with CAI and tAI - only accounting for one of these factors still suggested significant differences between the simulated regions. Although we are not the first to note potential issues with metrics like CAI or tAI for intragenic CUB analysis [16], our results demonstrate these metrics are insufficient for intragenic CUB analysis when these regions have drastically different amino acid usage or *φ* distributions, resulting in incorrect biological interpretation.

This is not to say CUB plays no role in the secretion of specific proteins. For example, experimental evidence demonstrates codon optimization of the *E. coli* maltose binding protein’s (MBP) signal peptide results in a decrease in protein abundance. Evidence suggests this is due to increased targeting of the codon optimized MBP by proteases due to improper folding [52, 53]. However, CUB as a means to guide proper co-translational folding is not a phenomenon unique to proteins with a signal peptide [4, 29, 48]. Although inefficient codons might be crucial to the fold of certain secreted proteins, our results do not indicate this is any more or less so than nonsecretory genes.

Although we found no general difference in selection on codon usage between signal peptides and the 5’-ends, it is possible CUB differences exist between the chaperone-dependent and chaperone-independent mechanisms of the Sec pathway. Previous analyses revealed patterns consistent with a region of slower translation at the 5’-ends of transmembrane proteins, which are typically secreted via SRP in bacteria [26]. [10] found transmembrane proteins in *E. coli* have a higher frequency of “programmed pause sites,” areas of high ribosomal density downstream from Shine-Dalgarno-like sequences, near the 5’-end. This region of higher ribosomal density was not observed in periplasmic proteins, which are normally secreted via SecA/B [26, 43]. Notably, [25] challenged the assertion that Shine-Dalgarno-like sequences are responsible for inducing translational pauses in bacteria, concluding signals previously seen were an artifact of the method for assigning ribosome occupancy along the transcript. [28] also found a consistent trend of inefficient codons 35-40 codons downstream of the SRP-binding site in various yeasts species using a modified form of the tAI. Ribosomal profiling data taken from *S. cerevisiae* provided experimental support for this hypothesis, but this analysis was limited to a small, closely-related phylogeny. Further work is needed to determine the general mechanistic role, if any, of codon-induced inefficient translation in SRP-dependent protein secretion, as well as to determine if any specific codon biases exists for SecA/B-dependent or chaperone-independent secreted proteins.

We do find selection on CUB is weaker at the 5’-ends relative to later portions of the gene, corroborating previous work [9, 13, 11, 16, 32, 35]. Weaker selection at the 5’-ends is often attributed to selection against nonsense errors and selection against mRNA secondary structure. Importantly, the advent of ribosome profiling suggested the presence of high ribosomal density at the 5’-ends, often referred to as the “5’-ramp” [44]. The 5’-ramp was originally thought to be the result of increased selection for slow translation at the 5’-end to reduce ribosomal interference further down the transcript, but simulations suggest the 5’-ramp is an artifact of short genes with high initiation rates [39]. Selection for co-translational folding is also thought to shape intragenic CUB [4, 29, 48]. Further work is needed to understand how these various selective forces are balanced to maintain translation efficiency and efficacious protein biogenesis.

Although it may be tempting to explain statistically significant results in the context of selection and adaptation, it is important to assert results cannot be explained by nonadaptive evolutionary forces (e.g. mutation bias and genetic drift) and/or as an artifact of some other constraint on the trait of interest (e.g. amino acid biases). We are certainly not the first to note the importance of considering nonadaptive explanations. Almost four decades ago, [14] critiqued the propensity of evolutionary biologists to invoke natural selection and adaptation without seriously considering possible nonadaptive explanations. The explosion of genomic data means now, more than ever, biologists should be hesitant to adopt adaptationist explanations to biological phenomenon without first investigating if such results could be shaped by nonadaptive forces. The embrace of “big data” by biological researchers is a double-edged sword: while we have the ability to investigate patterns and explore hypotheses which would not have been possible 20 years ago, the indiscriminate analysis of large datasets can lead to spurious, but statistically significant p-values, which are often misinterpreted as both evidence of a strong effect and a small probability of the null hypothesis being true. The misinterpretation of p-values and a bias towards adaptationist explanations can be a dangerous combination, leading to a misinterpretation of results and, in turn, misleading other researchers.

The development of models incorporating both adaptive and nonadaptive evolutionary forces will be important for understanding the selective forces shaping complex biological data. In the case of the studying CUB, codon indices like CAI have long been employed, but these metrics often are sensitive to and, thus, unable to disentangle the effects of amino acid and mutation biases from selection. While often good proxies of gene expression, these indices do not directly incorporate gene expression information into the weights estimated for each codon. This could lead to further problems of conflating mutation bias with selection when comparing CUB across regions. In contrast, because ROC-SEMPPR is grounded in population genetics and thus, is able to decouple selection and mutation bias, it serves as a more accurate and evolutionarily-grounded tool for the study of CUB. Ultimately, our work further illustrates the value of employing population genetics models which include nonadaptive evolutionary forces for analyzing genomic data.

## Acknowledgments

The authors acknowledge financial support from NSF grant MCB-1546402 (Primary Investigator: A. von Arnim), NSF grant MCB-1120370 (Primary Investigator: M.A. Gilchrist), the U.S. Department of Energy, Office of Science, and the Graduate School of Genome Science and Technology (University of Tennessee, Knoxville). Additional support was provided by the National Institute for Mathematical and Biological Synthesis (NSF:DBI-1300426 with additional support from the University of Tennessee).

